# Tuning Minimum-Norm regularization parameters for optimal MEG connectivity estimation

**DOI:** 10.1101/2023.04.15.537017

**Authors:** Elisabetta Vallarino, Ana Sofia Hincapié, Karim Jerbi, Richard Leahy, Annalisa Pascarella, Alberto Sorrentino, Sara Sommariva

**Affiliations:** Dipartimento di Matematica, Università di Genova, Genova, Italy; Computational and Cognitive Neuroscience Lab, Psychology Department, Université de Montréal, Montréal, Québec, Canada; MEG Center, Psychology Department, Université de Montréal, Montréal, Québec, Canada; MILA (Quebec Artificial Intelligence Institute), Montréal, Québec, Canada; Unique Center (Québec Neuro-AI Research Center), Montréal, Québec, Canada; Ming Hsieh Department of Electrical and Computer Engineering, University of Southern California, Los Angeles, California, USA; Istituto per le Applicazioni del Calcolo, Consiglio Nazionale delle Ricerche, Roma, Italy

**Keywords:** Functional connectivity, MEG, surrogate data, regularization parameter, minimum norm estimate

## Abstract

The accurate characterization of cortical functional connectivity from Magnetoencephalography (MEG) data remains a challenging problem due to the subjective nature of the analysis, which requires several decisions at each step of the analysis pipeline, such as the choice of a source estimation algorithm, a connectivity metric and a cortical parcellation, to name but a few. Recent studies have emphasized the importance of selecting the regularization parameter in minimum norm estimates with caution, as variations in its value can result in significant differences in connectivity estimates. In particular, the amount of regularization that is optimal for MEG source estimation can actually be suboptimal for coherence-based MEG connectivity analysis. In this study, we expand upon previous work by examining a broader range of commonly used connectivity metrics, including the imaginary part of coherence, corrected imaginary part of Phase Locking Value, and weighted Phase Lag Index, within a larger and more realistic simulation scenario. Our results show that the best estimate of connectivity is achieved using a regularization parameter that is 1 or 2 orders of magnitude smaller than the one that yields the best source estimation. This remarkable difference may imply that previous work assessing source-space connectivity using minimum-norm may have benefited from using less regularization, as this may have helped reduce false positives. Importantly, we provide the code for MEG data simulation and analysis, offering the research community a valuable open source tool for informed selections of the regularization parameter when using minimum-norm for source space connectivity analyses.

**Highlights:**

- The regularization parameter of the Minimum Norm Estimate of neural activity impacts connectivity estimation
- We study empirically the optimal parameter for connectivity estimation using realistic synthetic datasets
- We find the optimal parameter for connectivity estimation is systematically smaller than the optimal parameter for source imaging; different connectivity metrics yield the same result
- Code and data are available open source.

## 1. Introduction

During the past 20 years, examining functional connectivity has become a popular method to explore how brain networks behave in both healthy and diseased conditions, expanding the field of human brain mapping beyond analyzing local functions to encompass whole-brain functional networks (Singer, 1999; Varela et al., 2001; Friston, 2011; Sporns, 2012; Maestú et al., 2019). The fundamental concept is to identify and measure the degree of interactions among spatially distinct brain regions by analyzing the statistical correlation between their time courses (Fries, 2005; Pereda et al., 2005; Sakkalis, 2011; O’Neill et al., 2015).

Out of the various brain imaging methods, Magnetoencephalography (MEG) offers an interesting combination of excellent temporal and spatial resolutions, making it a promising tool to advance our understanding of functional brain connectivity in humans (Hämäläinen et al., 1993; Dalal et al., 2009; Schölvinck et al., 2013; Niso et al., 2022). MEG recordings offer a high temporal resolution, typically in the millisecond range, enabling the investigation of high-frequency brain dynamics that are beyond the reach of other imaging modalities, such as functional Magnetic Resonance Imaging (fMRI). The richness of the MEG signals in terms of spatial, temporal, and spectral information, along with the intricacy of the source reconstruction methods, can make MEG data more challenging to analyze and interpret compared to other types of neuroimaging signals. Although MEG signals can be analyzed at the sensor level, source-level analyses allow for more interesting anatomically grounded interpretations, and can arguably alleviate to some extent ambiguities caused by magnetic field spread (equivalent to volume conduc-tion in EEG) which makes interpreting sensor-level connectivity results more challenging.

Instead of investigating connectivity in sensor space, a more informative strategy is to solve the inverse problem (i.e., estimate the underlying current sources) and then assess the coupling between pairs of time series in the source space, which typically consists of a large number (e.g. 10, 000) of voxels in a brain volume (or nodes in a cortically constrained surface). While most studies of source-space connectivity employ this two-step approach (Schoffelen and Gross, 2019), there are few noteworthy exceptions (Fukushima et al., 2015; Ossadtchi et al., 2018; Tronarp et al., 2018). Although source-space connectivity is often preferred over sensor-level connectivity computations, the elaborate analysis pipeline it requires involves a series of decisions by the user, such as the choice of a source estimation algorithm, a connectivity metric and a cortical parcellation, to name but a few. While useful guidelines have been proposed (Gross et al., 2013; Pernet et al., 2020; Niso et al., 2022), certain decisions regarding the choice of methods and parameters still involve subjective judgment and fine-tuning through trial and error. Grasping the downstream consequences of methodological choices made at the beginning of a complex pipeline that goes from raw sensor data all the way to source space connectivity, is non-trivial.

One prominent choice is the selection of a parameter known as the *regularization* parameter. Regularization is necessary for MEG source reconstruction due to the ill-posed nature of the inverse problem of estimating neural source time-courses from MEG sensor data. The resulting solution is not unique, as many source configurations can explain the data equally well, and it is not continuously dependent on the data due to small perturbations from noise leading to large variations in the solution (Fokas et al., 2004; Dassios et al., 2005; Sarvas, 1987). Regularization is a classical mathematical approach to mitigate the ill-posedness in inverse problems, where uniqueness is restored by incorporating prior information on the solution (Hanke and Hansen, 1993; Baillet et al., 2001; Sorrentino and Piana, 2017; Ilmoniemi and Sarvas, 2019). Tikhonov regularization is a typical regularization method used in Minimum Norm Estimation (MNE), a widely used MEG inverse solver (Hämäläinen and Ilmoniemi, 1994), which consists in selecting among all possible solutions the one with minimum power. Over the past 10 years there have been over 1300 papers that have used or discussed the use of minimum-norm estimation for MEG analysis (query “Minimum norm” AND MEG performed on PubMed Central^®^ at the time of writ-ing). Within MNE, the weight assigned to prior information can be tuned by the regularization parameter. While it is known that too much regularization yields solutions that are too smooth, selecting a parameter that provides just the right amount of regularization often entails some degree of subjectivity. Furthermore, while one might intuitively expect that the amount of regularization that optimizes the estimation of MEG source time series would also optimize source space connectivity, evidence from previous simulated data suggests that this is not the case. In the first empirical evidence of this fact (Hincapié et al., 2016), the authors showed that the regularization parameter providing the best source-level coherence estimate is two orders of magnitude lower than the one providing the best estimate of source spectral power distribution. In the more recent mathematical treatment of the problem Vallarino et al. (2020) formally proved that the optimal regularization parameter for reconstruction of the cross-power spectrum, a largely used intermediate step in the connectivity estimation, is smaller than the optimal regularization parameter for source reconstruction; however, formal treatment required the strong assumption that neural signals and noise were white-noise Gaussian processes. In a numerical follow-up, the same authors showed that the inequality holds true when the data are generated by Multi-Variate Auto-Regressive models, and that the optimal regularization parameter for cross-spectrum estimation is related to a notion of spectrum complexity, or cross-spectrum signal-to-noise ratio (SNR) (Vallarino et al., 2021). However, these last two studies limited their analysis to the estimation of the cross-power spectrum, without considering the actual connectivity metric.

In this technical note, we address the question of determining the optimal regularization parameter for source-space connectivity detection, in a minimum-norm source estimation setting. The present study expands on previous work in three ways. Firstly, we investigate how the regularization parameter affects common state-of-the-art connectivity metrics, rather than the cross-power spectrum. Secondly, we conduct more extensive and diverse simulations with over 100, 000 distinct MEG cortico-cortical coupling configurations. Thirdly, we not only present the results of the numerical simulation, but also provide the pipeline and open source Python code in the hope to encourage other researchers to reproduce and extend our analyses. The code used to generate the data, example code, and codes for reproducing the paper’s plots are all available at https://themidagroup.github.io/regconnectivity/.

## 2. Methods and Materials

### 2.1 MEG data simulation

Sensor level MEG recordings were simulated by exploiting the linear model obtained from Maxwell’s equations under the quasi-static hypothesis (Hämäläinen and Ilmoniemi, 1994), i.e

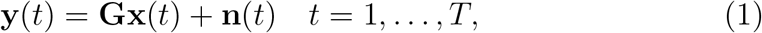

where at each time *t*, **y**(*t*) ∈ ℝ^*M*^, **x**(*t*) ∈ ℝ^*N*^ and **n**(*t*) ∈ ℝ^*M*^ represent the recordings at *M* sensors, the brain activity in *N* cortical locations, and the measurement noise, respectively, while **G** is the leadfield matrix. We used a leadfied matrix available within the MNE Python package (Gramfort et al., 2014). Following the recommendations in MNE-Python documentation, the available source space, containing 312, 273 sources, was down-sampled to 6940 dipole locations. Additionally, we only considered magnetometers and set the source orientations to be normal to the local cortical surface, so that **G** ∈ ℝ^*M×N*^ where *M* = 102 and *N* = 6940, and each column **G**_*i*_, *i* = 1, …, *N*, represents the magnetic field generated by a unit dipole placed at the *i*-th point of the source-space. We simulated *T* = 10, 000 time points to mimic about 78sec of recordings sampled at 128Hz.

MEG recordings were generated with the following pipeline structured in six steps (**S1**–**S6**) and schematically depicted in figure 1A.

**Figure 1:**
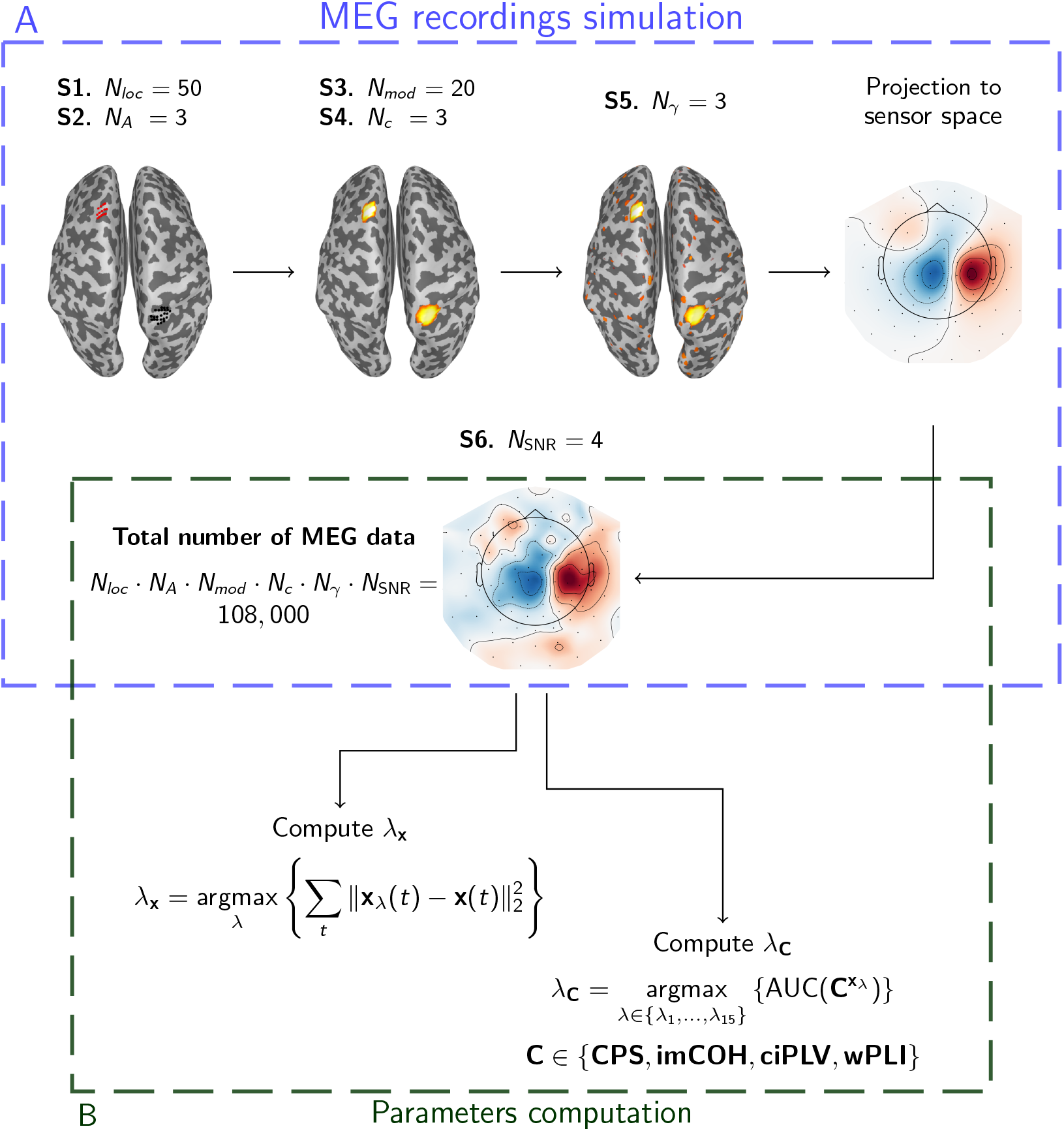
Pipeline of the MEG recordings simulation (A) and parameters computation (B). First, *N*_*loc*_ = 50 pairs of patches with *N*_*A*_ = 3 different extensions were generated (**S1** and **S2**). Then, *N*_*mod*_ = 20 MVAR models were exploited to generate the time series associated with the patches, each of them varying according to *N*_*c*_ = 3 intra-patch coherence levels (**S3** and **S4**). Finally, biological background noise with *N*_*γ*_ = 3 different levels of intensity was added (**S5**), and the generated neural activity was projected to sensor level and perturbed with measurement noise having *N*_SNR_ = 4 different levels of SNR (**S6**). A total of 108, 000 MEG recordings was thus generated by combining all features. Functional connectivity was estimated from each of the simulated MEG data through a two-step procedure consisting in estimating the neural activity and computing the corresponding connectivity metrics. The generated MEG recordings were then used to compute the optimal parameters for estimating neural activity (*λ*_**x**_) and connectivity (*λ*_**C**_, with **C** ∈ {**CPS, imCOH, ciPLV, wPLI**}).

#### S1. Set source locations

We randomly drew *N*_*loc*_ = 50 different pairs of source locations (patch–centers) so that their distance is greater than 4 cm and that their intensity at sensor space level is similar. To this end, we impose the ratio of the norm of the corresponding leadfield columns to range from 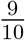 to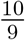.

#### S2. Set patches extension

For each pair of patch–centers we generated *N*_*A*_ = 3 different pair of patches by varying their extension, which was set equal to *A* = 2 cm^2^, 4 cm^2^, 8 cm^2^, similarly to what was done by Hincapie et al. (2017).

#### S3. Generate patch–centers time series

We generated *N*_*mod*_ = 20 different pairs of time series from as many MVAR models of order *P* = 5 with directional coupling from the first time series to the second (Haufe and Ewald, 2019; Sommariva et al., 2019), i.e.

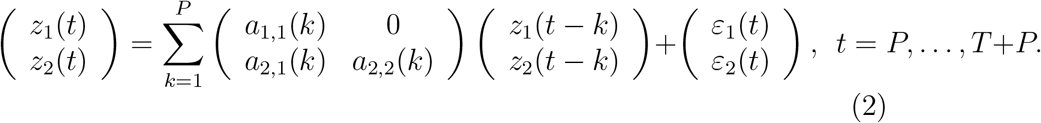

The non-zero elements *a*_*i,j*_(*k*) of the coefficient matrices were drawn from a normal distribution of zero mean and standard deviation Γ, whose value was randomly drawn in the interval [0.1, 1] so to obtain time series with different spectral properties (Vallarino et al., 2021). We retained only coefficient matrices providing (i) a stable MVAR model (Lütkepohl, 2005) and (ii) pairs of signals (*z*_1_(*t*), *z*_2_(*t*))^⊤^ such that the *ℓ*_2_-norm of the strongest one was less than 3 times the *ℓ*_2_-norm of the weakest one. The resulting time series (*z*_1_(*t*), *z*_2_(*t*))^⊤^ were normalised by the mean of their standard deviations over time, so that pairs of time series drawn from different models had similar magnitude. Finally, the obtained time series were filtered in the alpha range, i.e. in 8 − 12 Hz. For each pair of patches defined in **S2**, the activity of their centers was set according to the resulting time series.

#### S4. Generate patches of activity

The activity of each point of a patch was simulated by modulating the time courses of its center through a Gaussian window so that source magnitude decreased for increasing distance from the central source. The Fourier transform of the resulting time series were then perturbed in order to obtain *N*_*c*_ = 3 different levels of intra-patch coherence, namely *c* = 0.2, 0.5, 1. To this end, similarly to (Hincapie et al., 2017), we added at each frequency of the Fourier transformed data a random value drawn from a normal distribution with zero mean and random variance tuned so to obtained the desired level of coherence.

#### S5. Define background biological noise

To simulate the presence of background brain activity, each point of the source-space that did not belong to any patch was assigned with a time series following a univariate AR model of order 5. The overall resulting background activity was then scaled so that the inverse signal-to-noise ratio (SNR) between background activity and patches activity assumed *N*_*γ*_ = 3 different values, namely *γ* = 0.1, 0.5, 0.9 (Haufe and Ewald, 2019). Here SNR was defined as

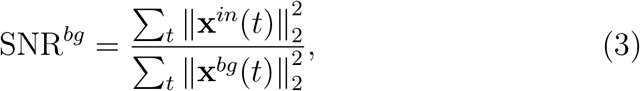

where **x**^*in*^(*t*) is the activity of interest, i.e the patches activity, and **x**^*bg*^(*t*) is the background activity.

#### S6. Define measurement noise

Finally, the simulated brain activity was projected to sensor level by means of the leadfield matrix and white Gaussian noise was added to obtain *N*_SNR_ = 4 different levels of SNR evenly spaced between -20 dB and 5 dB. In this case SNR was defined as

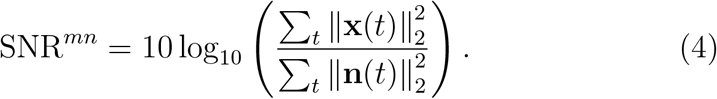

By combining all the mentioned features we obtained *N*_*loc*_ *· N*_*A*_ *· N*_*mod*_ *· N*_*c*_ *· N*_*γ*_ *· N*_SNR_ = 108, 000 different sensor level configurations.

### 2.2 MEG data analysis pipeline

Estimates of neural activity and connectivity were obtained from the simulated MEG data through the following standard two-step approach (Schoffelen and Gross, 2019).

i. First, a regularised estimate, **x**_*λ*_(*t*), of the neural activity, **x**(*t*), is obtained by solving the MEG inverse problem associated with (1). Here we consider as inverse solution the Minimum Norm Estimate (MNE) (Hämäläinen and Ilmoniemi, 1994), which is defined as

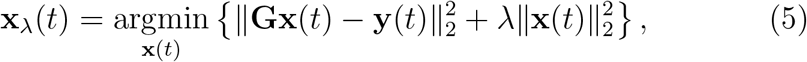

where *λ* is a proper regularisation parameter and ∥ *·* ∥_2_ is the *ℓ*_2_-norm.
ii. Then, a proper connectivity metric, 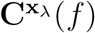, is computed from the estimated time series, **x**_*λ*_(*t*). Specifically, 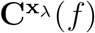 is a one parameter family of matrices of dimension *N × N*, whose (*j, k*)-th entry, 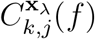, quan-tifies the interaction between two sources placed at the *j*-th and *k*-th location of the source space.

In this work, we considered four different connectivity metrics, namely the cross-power spectrum (CPS), the imaginary part of Coherence (imCOH) (Nolte et al., 2004), the corrected imaginary part of Phase Locking Value (ciPLV) (Bruña et al., 2018), and the weighted Phase Lag Index (wPLI) (Vinck et al., 2011). CPS was chosen in order to benchmark our results with those from previous works (Vallarino et al., 2020, 2021), while imCOH, ciPLV and wPLI were chosen as they are among the most widely used connectivity metrics and have been specifically designed to mitigate the effects of volume conduction or field spread by ignoring instantaneous zero-phase interactions. For all these connectivity metrics we used the implementations available in the MNE-python package (Gramfort et al., 2013). A concise mathematical formulation of the considered metrics can be found in Appendix A, and for further details we refer the reader to the mentioned papers.

### 2.3 Optimality criteria for choosing the regularization parameter

When the two-step approach described in the previous section is used to infer functional connectivity, the regularization parameter *λ* in (5) needs to be set to solve the MEG inverse problem in step (i). Previous studies (Hincapié et al., 2016; Vallarino et al., 2020, 2021) have shown that the value of *λ* needs to be set differently depending on whether one wants to obtain the best possible estimate of local neural activity or the best possible connectivity estimate. In other words, the value of *λ* that yields the best source reconstruction does not by extension lead to the best results when it comes to estimating connectivity from the reconstructed time series. In the present work the best solutions in the two contexts are identified according to the following criteria.

Inspired by Vallarino et al. (2020), given the the MNE estimate in (5), we define the optimal regularisation parameter for the reconstruction of **x**(*t*) as

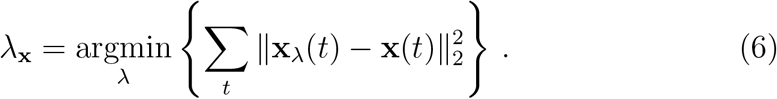

After computing *λ*_**x**_, for each of the four considered connectivity metrics, the corresponding optimal regularization parameter is defined as in Hincapié et al. (2016):

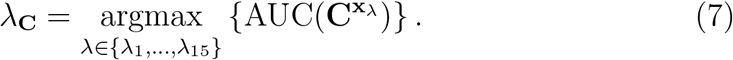

In the above equation {*λ*_1_, …, *λ*_15_} is a set of 15 possible regularization parameters obtained by multiplying *λ*_x_ for 15 values logarithmically spaced in the interval [10^−5^, 10^1^], and AUC stands for the Area Under the Receiver Operating Characteristic (ROC) curve, which is computed by plotting the True Positive Fraction (TPF) versus the False Positive Fraction (FPF) at different threshold levels.

We performed a seed-based connectivity analysis by considering interactions between the sources within the first patch (i.e. the patch associated with the time series *z*_1_(*t*) of (2)) and all other locations in the source-space. Hence, for a fixed threshold value *τ*, the TPF and FPF are defined as

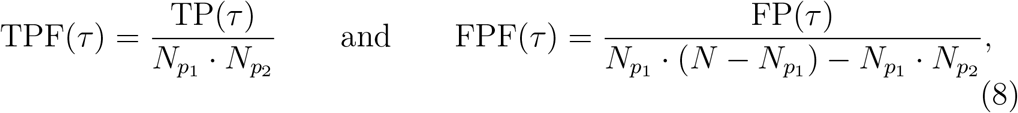

where 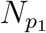 and 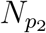 are the number of sources within the first and the second patches, respectively; TP(*τ*) represent the true positives (defined as the number of connections between sources belonging to the first patch and sources belonging to the second patch whose estimated intensities are above the threshold level *τ*); and FP(*τ*) represents the false positives (defined as the number of connections between sources belonging to the first patch and sources not belonging to any patch whose estimated intensities are above *τ*). Given a connectivity metric **C**, for each pair of sources the intensity of their connection is computed as

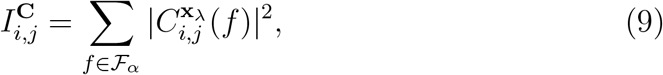

where ℱ_*α*_ is the alpha range 8–12Hz.

From a computational point of view, the minimization problem in (6) was solved through the Nelder-Mead simplex method (Nelder and Mead, 1965; Wright, 1996) implemented in SciPy (Virtanen et al., 2020), while *λ*_**C**_ was computed through custom code implemented in our labs.

In sum, based on the above definitions, we computed five different parameters; specifically, equation (6) defines the regulrization parameter providing the best possible estimate of the neural activity, while equation (7) is used to derive four optimal regularization parameters, one for each of the four considered connectivity metrics, i.e. **C** ∈ (**CPS, imCOH, ciPLV, wPLI**).

## 3. Results

### 3.1. Plot of the estimated source activity and connectivity for a representative case

To illustrate the typical output of our pipeline, figure 2 shows the results for one of the simulated MEG data. The original neural activity is depicted in the upper-right corner and consists of two active patches, one per hemisphere, where the patch in the right hemisphere leads the activity of the other one and will thus be considered as seed. As for the other parameters, in this simulation the area of the patches was *A* = 4 cm^2^, the level of intra-patch coherence was *c* = 0.5, the background biological noise had SNR^*bg*^ = 5 (i.e. *γ* = 0.2), and the measurement noise had SNR^*mn*^ = 5 dB. Since imCoh, ciPLV and wPLI are insensitive to zero-lag connections, the values of these connectivity metrics estimated between pair of sources belonging to the seed patch (i.e. the one in the right hemisphere for this simulation) resulted to be close to zero as shown in the bottom-right corner of figure 2. This does not apply to CPS which also accounts for the real part of the cross-spectrum.

**Figure 2:**
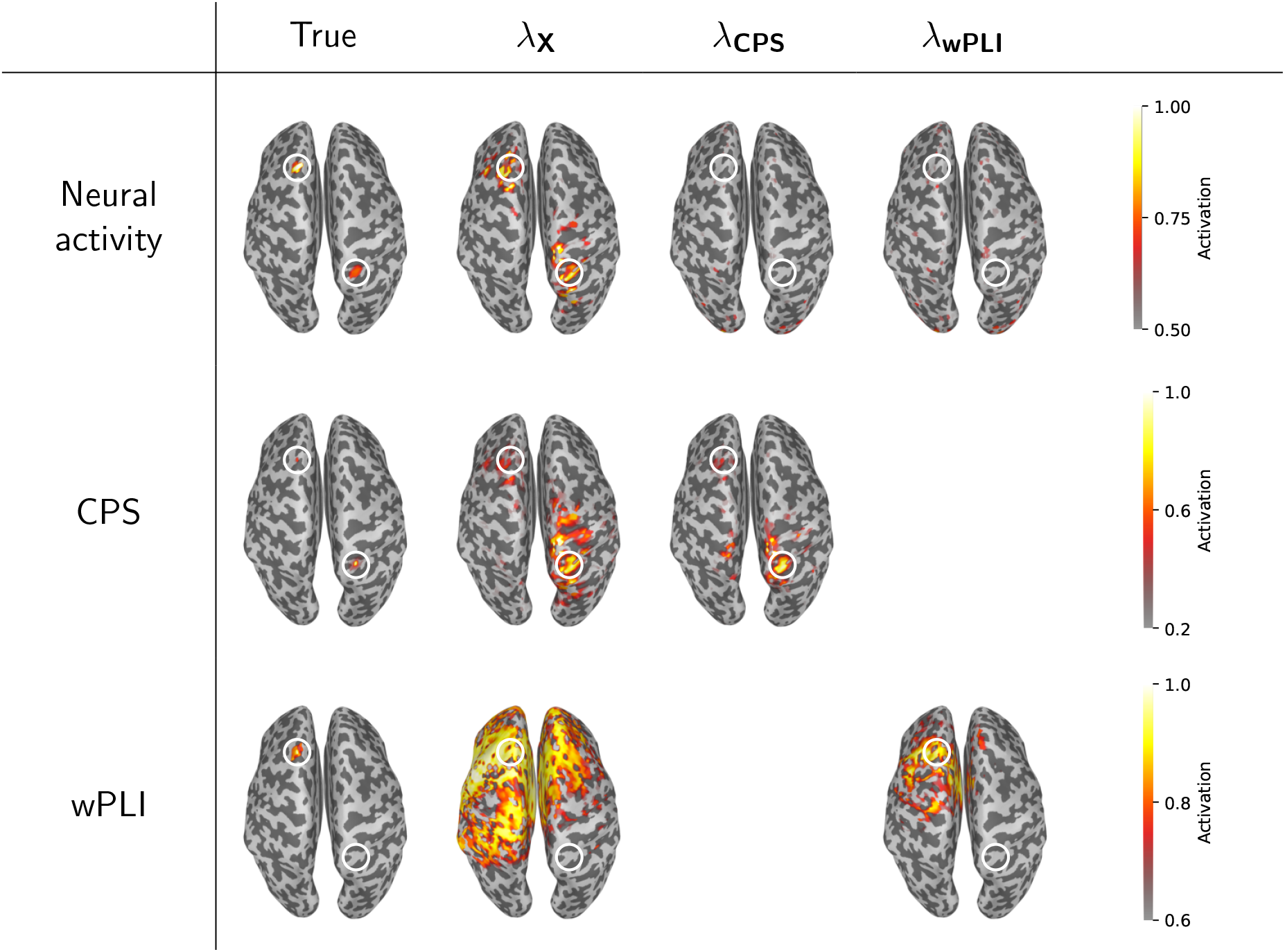
Neural activity and connectivity estimates for one simulated MEG data. The cortical maps in the different rows refer to neural activity, CPS and wPLI, respectively. The maps for the neural activity show the *ℓ*_2_-norm over time of the source time–courses. Since we performed a seed-based connectivity analysis, the maps for CPS and wPLI are obtained by averaging over the points of the seed–patch the values of the connectivity between such points and all points of the source-space. For each row, the first column shows the theoretical values computed from the simulated time-series, **x**(*t*), while the other columns show the estimates obtained by using the listed regularization parameters. White circles highlight the locations of the active patches. In this simulation we considered as seed-patch the one on the right hemisphere.

For the simulated configuration we then computed the optimal parameters for the estimation of neural activity, *λ*_**x**_, and connectivity, *λ*_**CPS**_, *λ*_**imCOH**_, *λ*_**ciPLV**_, *λ*_**wPLI**_, as described in the previous section. For this simulation *λ*_**x**_ = 123.68, while the parameter of all the connectivity metrics was about two order of magnitude lower, specifically *λ*_**CPS**_ = 0.4610, *λ*_**imCOH**_ = *λ*_**ciPLV**_ = *λ*_**wPLI**_ = 1.2360.

Figure 2 shows the benefits of using the appropriate regularization parameters. It is easy to see that the true values of neural activity, CPS and wPLI are reconstructed with higher accuracy when the corresponding optimal regularization parameters are used, i.e. *λ*_**x**_, *λ*_**CPS**_ and *λ*_**wPLI**_, respectively. Conversely, if either *λ*_**CPS**_ or *λ*_**wPLI**_ are used instead of *λ*_**x**_, the reconstructed neural activity results are more noisy. This is consistent with the fact that lower values imply less regularization. For the same reason, if *λ*_**x**_ is used instead of either *λ*_**CPS**_ and *λ*_**wPLI**_, the estimated connectivity presents a higher number of spurious connections due to the fact that the underlying neural activity estimates are overly smoothed. The results of this simulation also illustrate a known pitfall of the two-step approach for connectivity estimation Palva et al. (2018). Indeed, the reconstructed connectivity maps for CPS and wPLI show a certain amount of spurious connections between sources in the vicinity of true interacting sources, and the number of such spurious connections is only partially limited by choosing the proper regularization parameter. The observations concerning wPLI also hold for imCOH and ciPLV. The analogous plots for these two connectivity metrics can be found in Figure S1.

### 3.2. Connectivity estimation requires less regularization

Figure 3 shows the effect of the regularisation parameter on connectivity estimation in terms of the AUC values. Specifically, for each of the tested parameters, (i.e *λ*_1_, …, *λ*_15_ of (7)) we show the AUC value averaged across all simulated data. The maximum value of AUC for CPS is around 0.77 and it is reached when *λ* is about one order of magnitude lower than *λ*_**x**_ (i.e. *λ* ≃ 0.72 · 10^−1^*λ*_**x**_). For the other connectivity metrics the maximum value of AUC is reached for a lower value of the parameter, i.e. *λ* ≃ 2.7 · 10^−2^*λ*_**x**_, and it is slightly higher, i.e. 0.83 for imCOH and wPLI and 0.81 for ciPLV. Moreover, the trend of the AUC value as a function of *λ* is more sharpened for imCOH, ciPLV and wPLI which may indicate that for these connectivity metrics choosing a sub-optimal parameter is more damaging. This effect appears to be less prominent with CPS, where the evolution of the AUC value as a function of *λ* is smoother.

**Figure 3:**
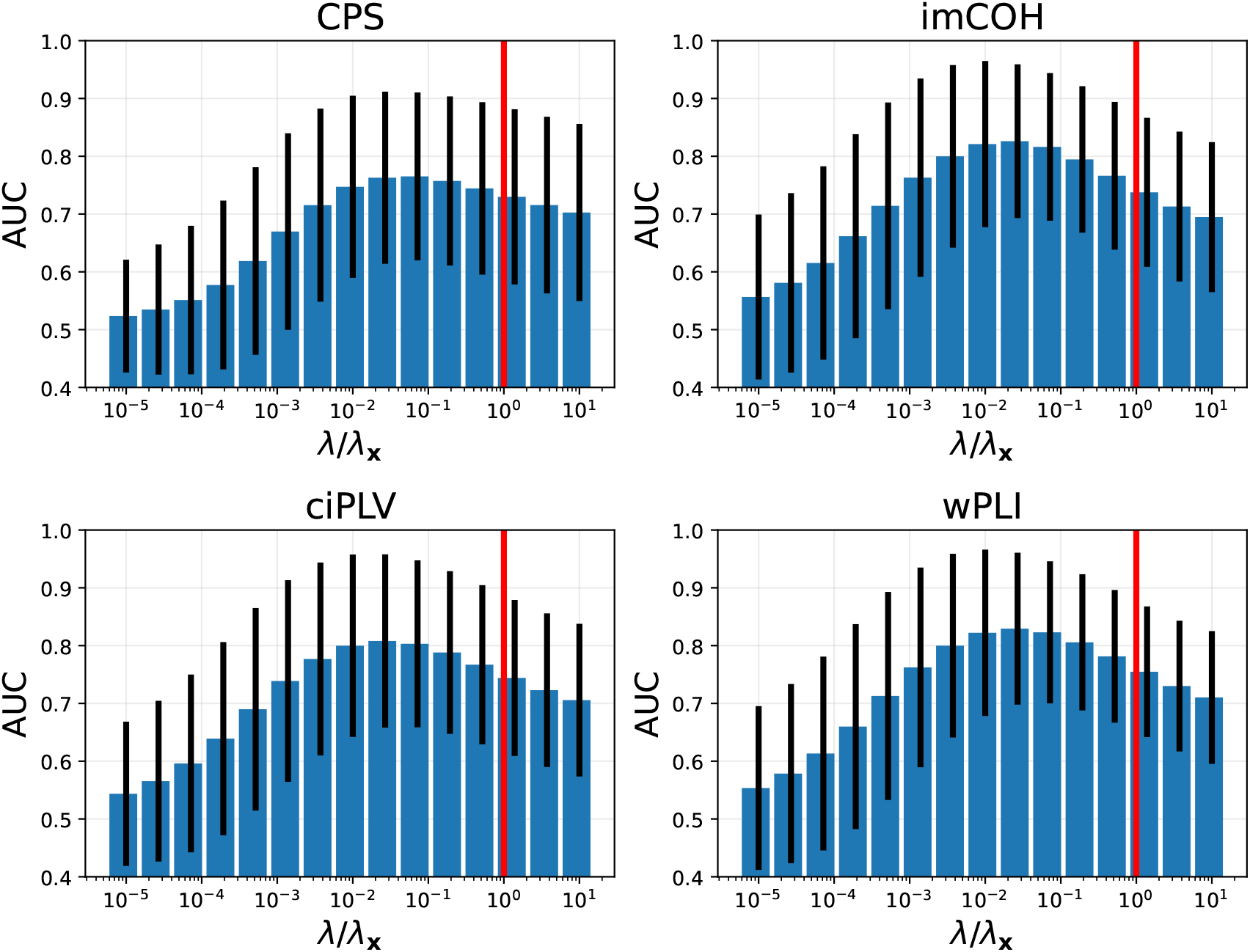
AUC values as function of the tested regularization parameter *λ* for the four connectivity metrics. In each panel, barplots and corresponding error bars represent mean and standard deviation of the AUC values across the 108, 000 simultations, while the x axis displays the value of the ratio between *λ* and the parameter *λ*_**x**_ providing the best estimate of the neural activity. The red vertical line highlight when *λ* = *λ*_**x**_

### 3.3. Different connectivity metrics behave similarly

Finally, we compared the optimal regularization parameters obtained across the different connectivity metrics, and investigated the impact of the different features of the MEG data, namely patch extension, intra-patch coherence, level of background noise and measurement noise.

In our simulations, the optimal regularization parameters for CPS, *λ*_**CPS**_, were on average lager than those for the other connectivity metrics, as can be seen from figure 4 (left panel). By contrast, imCOH, wPLI and ciPLV usually require a similar level of regularization as shown by figure 4 middle and right panel, where the 2D histograms lined up along the box diagonal.

**Figure 4:**
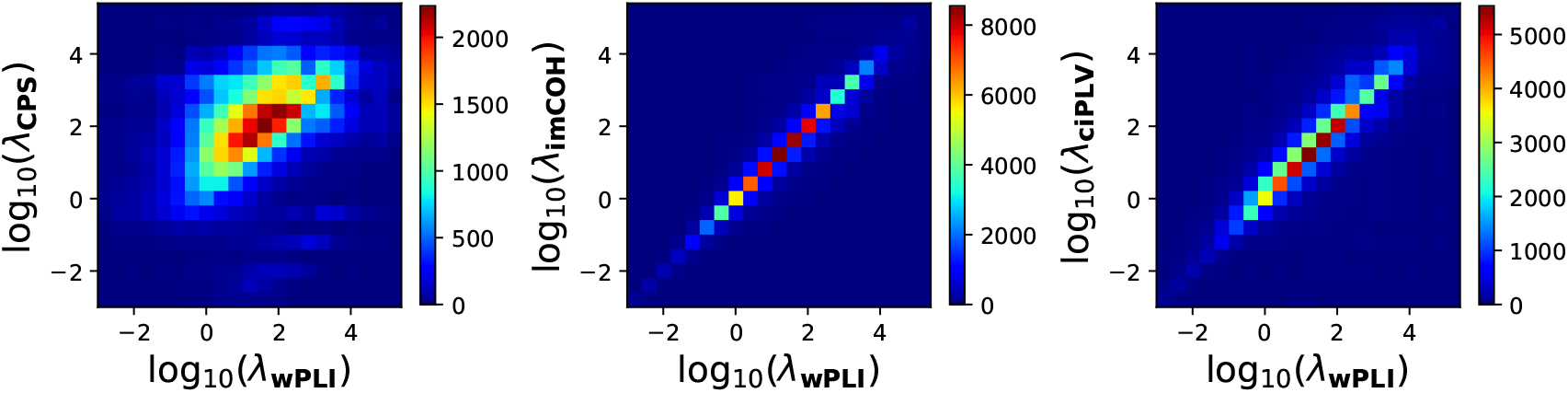
2D histogram showing the relationship between the optimal regularization parameters for different connectivity metrics. In each panel the x-axis shows the value of the optimal parameter for wPLI in logarithmic scale, while the y-axis refers to CPS, imCOH and ciPLV, respectively. Notice the different scale for the colorbar in each panel.

Interestingly, these observations did not dependent on the variations in patch extension, intra-patch coherence, level of background noise or measurement noise. As an example, figure 5 shows the relationship between the optimal regularization parameters for the different connectivity metrics as a function of the level of measurement noise. As expected the lower the SNR^*mn*^ the higher the required regularization. However, regardless the actual value of SNR^*mn*^, *λ*_**CPS**_ was generally larger than *λ*_**wPLI**_, while the values *λ*_**imCOH**_ and *λ*_**ciPLV**_ were close to that of *λ*_**wPLI**_ as was already observed in figure 4.

**Figure 5:**
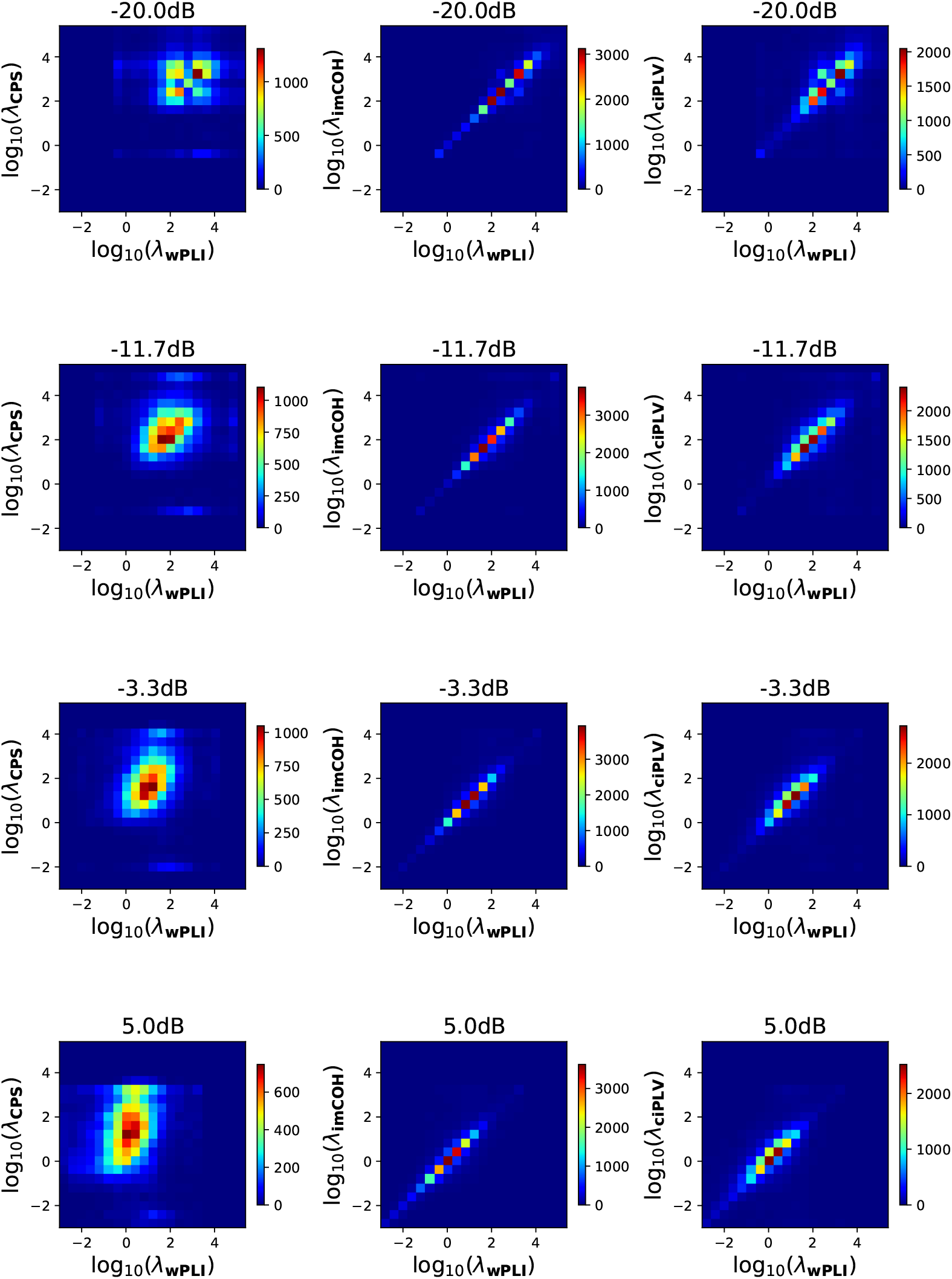
Impact of the measurement noise on the relationship between the optimal regularization parameters for different connectivity metrics. Each row refers to a different signal-to-noise ratio whose value is reported on top of the panels. Each column shows the 2D-histogram for a different pair of connectivity metrics, namely CPS vs wPLI (left column), imCOH vs wPLI (middle column), and ciPLV vs wPLI (right column). Each panel as been obtained similarly as in 4.

The results obtained with the other considered features were comparable (see supplementary figures S2, S3 and S4).

## 4. Discussion

The goal of the present technical note is to provide a thorough assessment of an often overlooked issue in MEG source-space connectivity work, which is the fact that achieving the best source localization and best inter-areal connectivity assessment requires different amounts of regularization. Although arguably intuitive, a systematic assessment of the relationship between the optimal regularization values for source localization and for connectivity detection has been missing. Using extensive simulations with a wide range of source configurations and noise conditions, we illustrate and examine this effect across multiple state-of-the-art connectivity metrics.

More specifically, we simulated a dataset of 108, 000 realistic MEG sensor recordings generated by coupled patches of cortical activities of variable size, location, and with different levels of intra-patch coherence, biological and measurement noise. For each simulated data set we computed five optimal regularization parameters which are respectively optimal for estimating the underlying neural activity, and estimating four different connectivity metrics, namely CPS, imCOH, wPLI and ciPLV.

Importantly, our simulations showed that the best estimate of connectivity is achieved using a regularization parameter that is roughly one to two orders of magnitude smaller than the one that yields the best source localization. This is a remarkable difference and may imply that previous work assessing source-space connectivity after using minimum-norm without adapting the regularization parameter for connectivity analysis, could in theory contain more false positives (i.e. spurious interactions) than expected. The present study extends the results of previous work by Hincapié et al. (2016) and Vallarino et al. (2020, 2021), in several ways, but especially by expanding the assessment to several popular state-of-the-art connectivity metrics that are routinely used by the MEG and EEG research community. In particular, we show here that the increased risk of detecting spurious interactions due to too much regularization is also relevant when using connectivity metrics that ignore zero-lag interactions. Our simulations show that the peak AUC for connectivity detection using imCOH, wPLI and ciPLV requires a regularization parameter that is 1-2 orders of magnitude smaller than the parameter that yields the peak AUC for source localization.

Overall, we found that imCOH, wPLI and ciPLV behaved similarly, which is coherent with the fact that they are all normalized connectivity metrics that are insensitive to zero-lag coherence. They also required a regularization parameter which was slightly lower than the optimal parameter for CPS. Furthermore, our simulations indicated that our findings concerning the various connectivity metrics we tested were robust to changes across a wide range of simulation features (i.e. patch extension, intra-patch coherence, SNR^*bg*^ and SNR^*mn*^).

Importantly, we provide all the code implemented for simulating the MEG recordings and for reproducing the plots of the paper. These are available at https://themidagroup.github.io/regconnectivity/ and, our hope is that this will provide a useful open source tool for the community to further examine the impact of regularization on source-space connectivity assessments using a wide range of metrics and coupling patterns.

From a pragmatic perspective, thinking about how our observations could translate into actionable recommendations, our study established the need for setting the regularization at 1-2 orders of magnitude lower than the optimal value for source localization, when the goal is to examine connectivity patterns. Doing so reduces the number of false positives. In real life situations, we obviously do not have access to the ground truth, and our analysis here does not provide a method to compute the optimal regularization parameter. A general recommendation would therefore be to use current best practices to determine the regularization parameter that is thought to optimize source reconstruction, and then scale it down by a factor of 10 to 100, if the aim is to compute source-space connectivity. Current data-driven approaches to setting the regularization parameter range from classical approaches based on regularization theory, such as L-curve method, generalized cross-validation, discrepancy principle (Hansen, 2005; Grech et al., 2008), to more recent learning techniques based on Bayesian hyper-parameter estimation and deep learning (Afkham et al., 2021). However these methods are seldom applied in MEG studies, where instead heuristic strategies linking the regularization parameter to the data SNR are often employed (Ramírez et al., 2011; Samuelsson et al., 2021).

As an alternative approach, the linearity of MEG forward problem and of the Fourier transform when computing CPS can be exploited to tackle the inverse problem by directly estimating source-space connectivity from scalp connectivity maps (Ossadtchi et al., 2018; Vallarino et al., 2020). Future work should examine how the findings reported here extend to regularization approaches used in other source estimation techniques, such as beamformers (Westner et al., 2022), as well as different generalizations of the minimum-norm approach.

Furthermore, in this study we only examined simulated MEG recordings. Future work should also investigate to which extent our findings hold true for other modalities, in particular when source-space connectivity is inferred from Electroencephalographic data with various sensors configurations (Allouch et al., 2022).

We hope that the simulations and illustrative examples provided in this technical note will help researchers make informed decisions and raise awareness about the importance of carefully choosing the regularization parameter in the context of source-space connectivity estimation. We also hope that the associated open source tools we made available will allow our research community to collaboratively expand on this work in an open science spirit.

//

## 5. Acknowledgments

K.J. is supported by funding from the Canada Research Chairs program (950-232368) and a Discovery Grant from the Natural Sciences and Engineering Research Council of Canada (2021-03426), a Strategic Research Clusters Program (2023-RS6-309472) from the Fonds de recherche du Québec - Nature and technologies.

E.V., A.P., A.S. and S.S. were partially supported by Gruppo Nazionale per il Calcolo Scientifico.

A.P. acknowledges the CNR for the Short Term Mobility 2019.

R.M.L. was supported by NIH R01 EB026299 from the National Institute of Biomedical Imaging and Bioengineering.

## Appendix A. Connectivity metrics

In this work we considered four connectivity metrics, namely the cross-power spectrum (CPS), the imaginary part of Coherence (imCOH) (Nolte et al., 2004), the corrected imaginary part of Phase Locking Value (ciPLV) (Bruña et al., 2018), and the weighted Phase Lag Index (wPLI) (Vinck et al., 2011). The definitions of such metrics follow below.

Suppose 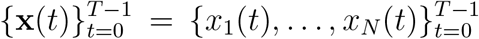 to be the time series associated with the activity of *N* sources in the brain along the time points *t* = 0, …, *T* − 1. CPS is defined by exploiting the Welch’s method (Welch, 1967), which consists in partitioning the data in *P* overlapping segments {**x**^*p*^(*t*)}_*p*=1,…,*P*_, computing the discrete Fourier transform of the signals multiplied by a window function *w*(*t*), 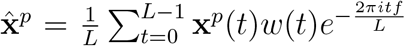, and then averaging these modified periodograms, i.e.

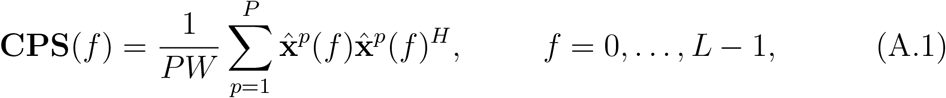

where *L* is the length of the segments and 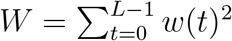.

Starting from CPS, imCOH between *x*_*j*_(*t*) and *x*_*k*_(*t*) is defined as

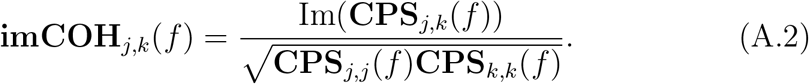

Finally ciPLV and wPLI are defined, respectively, as

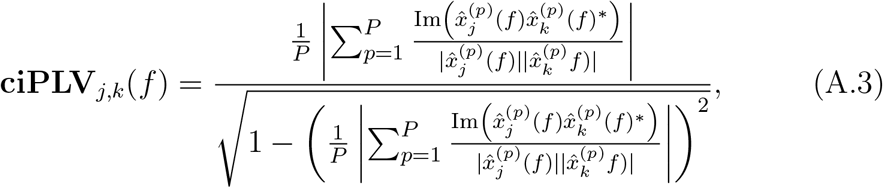

and

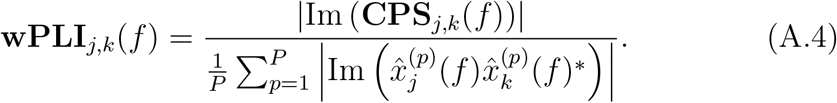

In all previous formula, Im(*z*) and *z*^*^ indicate the imaginary part and the complex conjugate of *z*, respectively.

## Notes

https://themidagroup.github.io/regconnectivity/

### Competing Interest Statement

The authors have declared no competing interest.

