## supplementary figures for "Tuning Minimum-Norm regularization parameters for optimal MEG connectivity estimation"

### Supplementary material for the manuscript Tuning Minimum-Norm regularization parameters for optimal MEG connectivity estimation

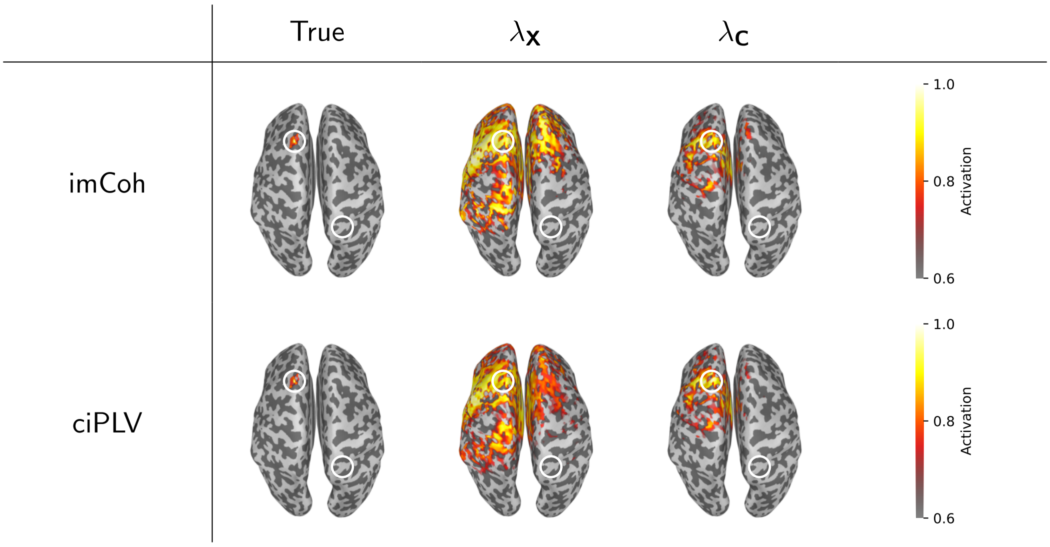


Figure S1: True and estimated connectivity cortical maps when imCOH (first raw) and ciPLV (second raw) are used. Similarly to Figure 2 of the manuscript, for each row, the first column shows the theoretical values computed from the simulated time-series, $\mathbf{x}(t)$, while the other columns show the estimates obtained by using the listed regularization parameters. White circles highlight the locations of the active patches. In this simulation we considered as seed-patch the one on the right hemisphere.

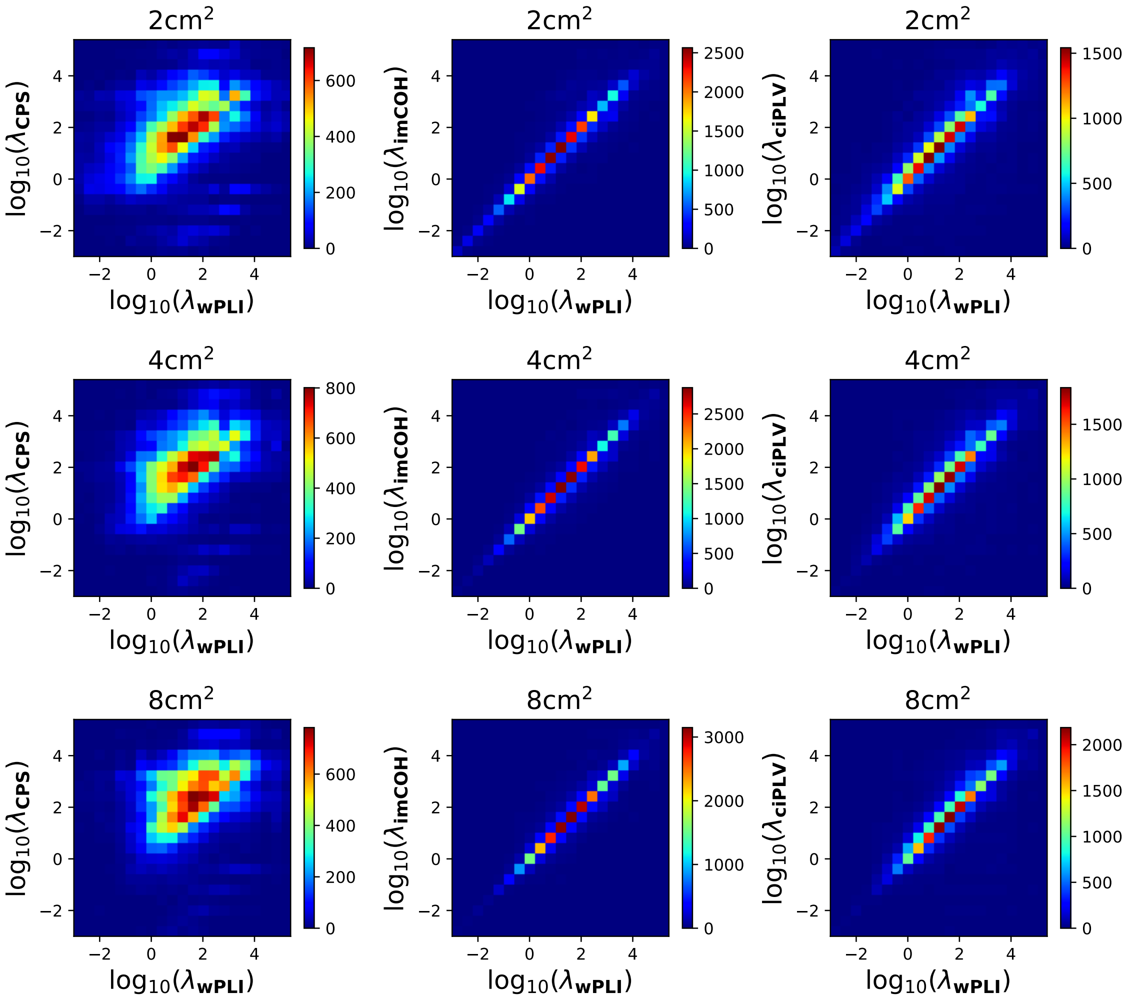


Figure S2: Impact of the patch extension on the relationship between the optimal regularization parameters for different connectivity metrics. Each row refers to a different extension of the active patches whose value is reported on top of the panels. The different columns and panels have been organized as in Figure 5 of the manuscript.

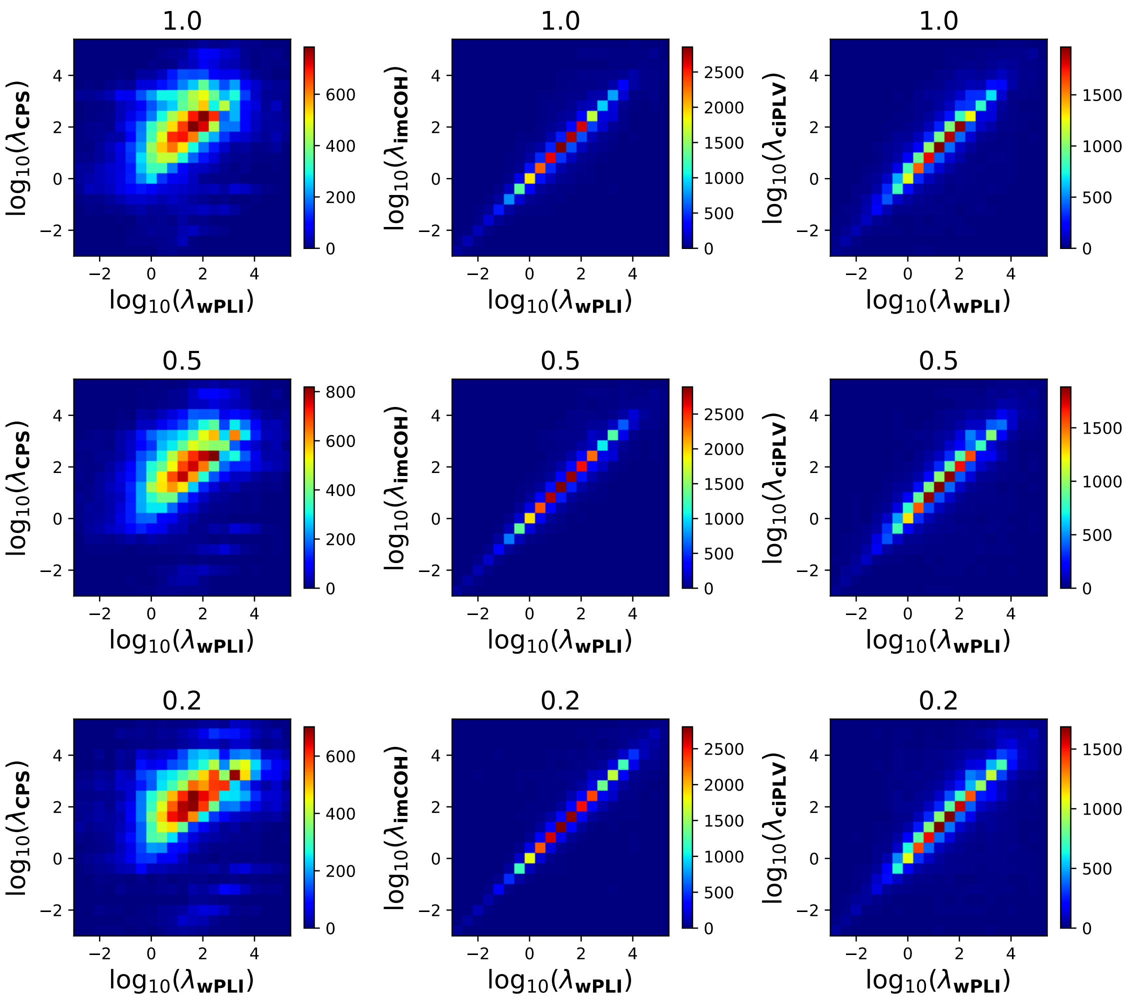


Figure S3: Impact of the intra-patch coherence on the relationship between the optimal regularization parameters for different connectivity metrics. Each row refers to a different level of intra-patch coherence whose value is reported on top of the panels. The different columns and panels have been organized as in Figure 5 of the manuscript.

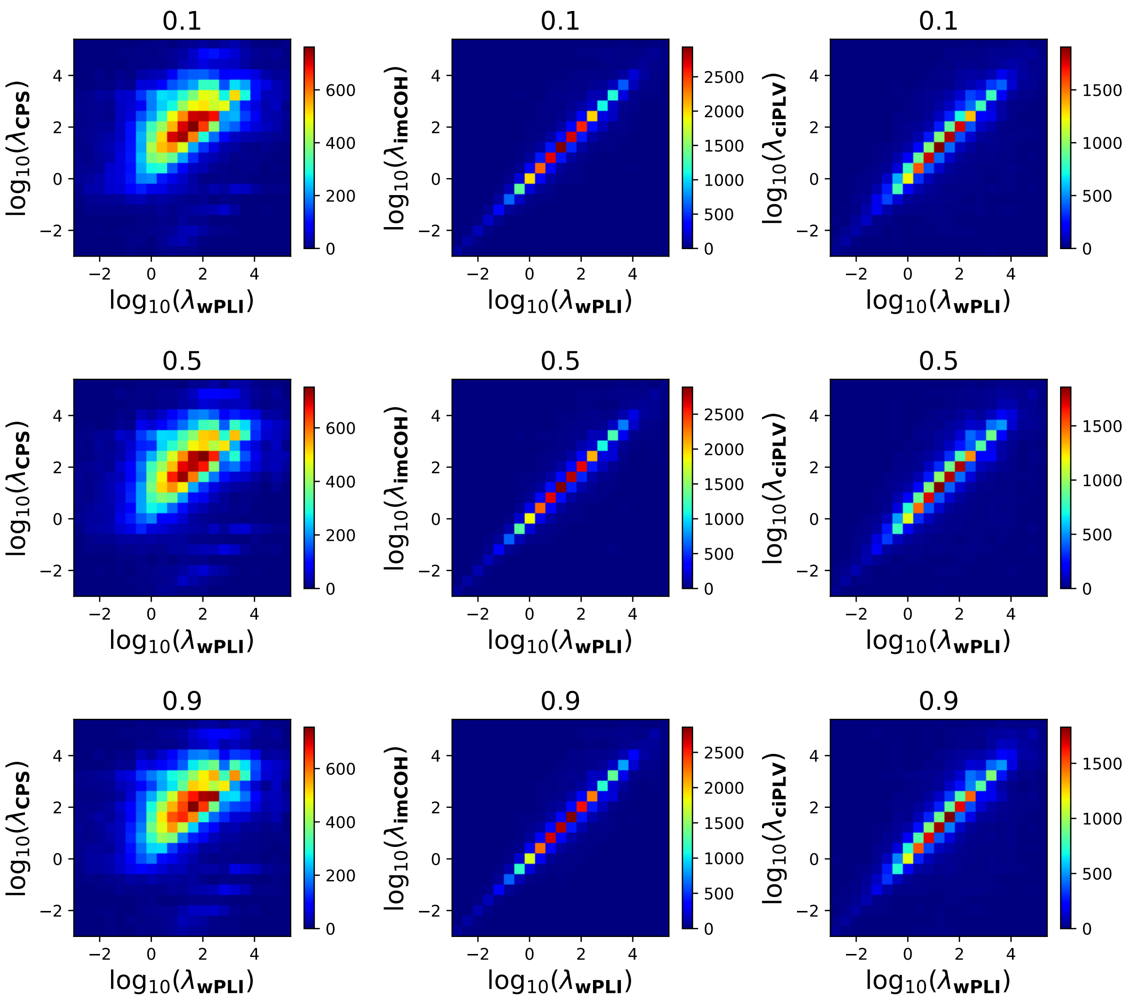


Figure S4: Impact of the background biological noise on the relationship between the optimal regularization parameters for different connectivity metrics. Each row refers to a different signal-to-noise ratio (SRN$^{bg}$) whose value is reported on top of the panels. The different columns and panels have been organized as in Figure 5 of the manuscript.
